# Motor effort biases perceptual decisions through later-stage evidence accumulation rather than early sensory-attentional processing

**DOI:** 10.64898/2026.10.05.756778

**Authors:** Ningjing Cui, Jeroen B. J. Smeets, Robert J. van Beers, Dock H. Duncan, Bernadette C. M. van Wijk

## Abstract

Motor effort has been shown to bias perceptual decisions toward less effortful choices, but it remains unknown whether this bias already takes effect during the early stages of decision formation through preparatory attentional allocation or sensory processing, or during later-stage evidence accumulation preceding response execution. This study investigated when motor effort becomes integrated into decision formation by focusing on the temporal evolution of neural processing during perceptual decision-making. While 64-channel EEG was recorded, twenty-four healthy adults performed an orientation discrimination task in which two Gabor patches appeared simultaneously in the left and right visual hemifield and decisions were reported using two pressure-sensitive response buttons requiring different levels of force. We found that motor effort modulated EEG activity at both the preparatory and later decision stages: pre-stimulus alpha power was reduced over the occipito-parietal cortex contralateral to the low-effort option, and mean centro-parietal positivity (CPP) amplitude preceding response execution was greater over centro-parietal cortex for low-effort choices, while leaving early sensory representations intact as indicated by no differences in P1 or N1 amplitude following stimulus onset. Trial-by-trial CPP amplitude preceding response execution increased with the likelihood of selecting the low-effort option, whereas the pre-stimulus alpha modulation was not associated with behavioral decision bias. These findings suggest that motor effort biases perceptual decisions primarily during later-stage evidence accumulation, thereby providing insight into the neural basis of embodied perceptual decision making.

## 1. Introduction

Perceptual decision making refers to the process of selecting one option from a set of alternatives based on available sensory evidence (Gold & Shadlen, 2007; Heekeren et al., 2008). In traditional lab-based settings, this process is commonly investigated using tasks such as random-dot motion discrimination (e.g., Hanks et al., 2006; Huk & Shadlen, 2005) or orientation discrimination paradigms (e.g., Assarioti et al., 2025; Zhang et al., 2019), in which decisions are reported through motor responses (e.g., button presses). Classical views conceptualize decision making as a serial process, in which sensory information is accumulated to form a choice before being translated into an action (Gold & Shadlen, 2007; Heekeren et al., 2008), with action execution considered merely as the final output of a decision process. However, in real-life scenarios, perceptual choices are embedded within our bodily actions and surrounding environments, and the actions associated with available options often differ in their costs and demands. For instance, when choosing between two equally desirable fruits located at different positions in an orchard, individuals are generally more inclined to select the fruit that requires less effort to reach. Such observations suggest that action costs can systematically bias choice behavior and have led to the development of the framework of embodied perceptual decision making (Cisek & Kalaska, 2010; Cisek & Pastor-Bernier, 2014; Lepora & Pezzulo, 2015), which proposes that perceptual decisions emerge through continuous coupling of sensory, cognitive, and motor processes, allowing the costs of potential actions to shape the process of choice formation.

The framework of embodied perceptual decision making has gained substantial support from behavioral studies in which the motor effort required to report different choices was manipulated independent of the perceptual evidence, such as through increased physical resistance (Burk et al., 2014; Hagura et al., 2017), greater reaching distance (Assarioti et al., 2025), or a more inconvenient movement angle (Marcos et al., 2015). These studies consistently showed that motor effort biases perceptual decisions toward less effortful options. Importantly, this motor-driven decision bias persists even when choices are reported vocally rather than through physical movement (Hagura et al., 2017), suggesting that it cannot be fully explained by transient response-selection tendencies alone, but may instead reflect a more global influence of motor-related information on perceptual decision formation. That is, motor contingencies may be integrated into the computations underlying perceptual decisions rather than acting solely at a downstream motor-output stage.

Beyond behavioral observations, converging neurophysiological evidence has further supported an active role of the motor system in perceptual decision making (Cisek & Kalaska, 2010; Cisek & Pastor-Bernier, 2014). At the neural level, sensory information is first encoded in visual cortical regions (Britten et al., 1996; Newsome & Pare, 1988) before being transmitted to parietal and frontal regions involved in evidence evaluation and choice formation (Heekeren et al., 2008; Huk & Shadlen, 2005; Platt & Glimcher, 1999; Zhou & Freedman, 2019). In non-human primates, neural activity within sensorimotor circuits has been shown to simultaneously represent multiple candidate actions and interact with upstream parietal and frontal choice-related regions (Cisek & Kalaska, 2005; Pastor-Bernier & Cisek, 2011), suggesting that the motor cortex participates in shaping how decisions unfold. Consistently, EEG and MEG studies in humans have demonstrated that sensorimotor beta oscillations over the motor cortex encode previous motor responses and bias subsequent perceptual decisions toward alternation (Cui et al., 2026; Pape & Siegel, 2016; Urai & Donner, 2022). More causal evidence comes from TMS studies showing that changes in corticospinal excitability within the motor cortex track the representation of competing choices in both perceptual (Michelet et al., 2010) and value-based decision tasks (Klein-Flügge & Bestmann, 2012).

Despite evidence that motor-related information may influence perceptual decisions, the processing stage at which this influence emerges remains unsolved. In other words, does the decision bias toward the lower-effort option when choosing between two equally desirable fruits at different positions stem from differences in how these options are visually perceived in the first place, or does the bias emerge only during later stages of decision formation? Addressing this question requires characterizing the temporal dynamics of the neural processes underlying perceptual decision-making. Here, we considered three candidate stages along the perception–action pathway at which motor-related information may influence perceptual decisions: preparatory attentional allocation preceding perceptual evidence onset, early sensory encoding of perceptual evidence, and later-stage evidence accumulation leading to choice commitment.

The first stage at which motor effort may influence decision formation is attentional allocation prior to the presentation of perceptual evidence. The finding that an asymmetry in action space due to an amputation yields a corresponding perceptual bias in position judgement in near visual space (Makin et al., 2010) can be interpreted as support for this option. One neural marker of such preparatory attentional allocation is alpha-band activity over the parieto-occipital cortex. Alpha oscillations (8–12 Hz) are prominent EEG rhythms observed bilaterally when attention is distributed across both visual hemifields, and spontaneous fluctuations in alpha activity are considered to reflect the momentary state of visual cortex excitability (Romei et al., 2008). Besides, a lateralized decrease in alpha power (i.e., alpha desynchronization) is thought to index enhanced spatial attention toward the contralateral visual hemifield (Kelly et al., 2006; Thut et al., 2006; Worden et al., 2000). Therefore, if motor effort alters pre-stimulus attentional states during perceptual decisions, a reduction in alpha power should be evident over the hemisphere contralateral to the more attended option.

A subsequent stage at which motor effort may influence decision formation is early sensory encoding, when the available options are first visually presented. If motor-related information is integrated early in the perception–action pathway, knowledge of the motor costs associated with the available options may alter the encoding of perceptual information itself, leading to differential sensory representations of the two options. Potential neural markers for such modulation are P1 and N1 components over the occipital cortex. The P1 and N1 are early visual event-related potentials (ERPs) occurring approximately 80–130 ms and 150–200 ms after stimulus onset, respectively, that are generated contralateral to the visual hemifield and have been associated with the early processing of visual information (Hillyard et al., 1998). In particular, the P1 has been associated with early sensory gain (Di Russo et al., 2002; Hillyard et al., 1998), whereas the N1 is considered to index selective discrimination of visual stimuli (Hopf et al., 2002; Vogel & Luck, 2000). Therefore, if motor effort alters the early sensory representation of the available options, P1 and N1 amplitudes may differ between the hemispheres contralateral to the two visual hemifields associated with the available options.

Lastly, prior to choice commitment, sensory evidence associated with options requiring different levels of motor effort may be accumulated differently, such that evidence favoring one option is integrated more efficiently than evidence favoring the alternative. A neural index of this evidence accumulation process is the centro-parietal positivity (CPP), an event-related potential measured over the centro-parietal cortex that gradually ramps up and reaches a plateau immediately before response execution (O’Connell et al., 2012). The CPP has been shown to track the accumulation of decision evidence, with greater amplitude or steeper buildup observed when sensory evidence is stronger or decisions are easier (Kelly & O’Connell, 2013; Twomey et al., 2015). Therefore, if motor effort biases decisions through evidence accumulation, differences in CPP amplitude may be evident prior to response execution when choosing between the options with different levels of required effort.

Prior work investigating behavioural responses with computational modeling indicated that motor effort might influence central decision processes rather than the early extraction and encoding of sensory input, as the observed bias was best accounted for by a static shift at the starting point of evidence accumulation (Hagura et al., 2017). This points to the possibility that motor-driven decision bias enters the decision process before evidence accumulation as a static bias towards one of the options, rather than by shifting the evidence accumulation process itself toward the easier option. Several other attempts have been made to investigate decision biases arising from non-motor sources such as prior probability, reward, and experimentally induced perturbations of neural activity. Most of those studies have similarly revealed that decision biases primarily arise from an initial bias prior to evidence accumulation rather than from changes in the evidence accumulation process (Ding & Gold, 2012; Hanks et al., 2006; Mulder et al., 2012), although effects on evidence accumulation were also observed when the reliability of the incoming sensory evidence was unpredictive across trials (Hanks et al., 2011). Computational modeling alone, however, cannot distinguish the neural processing stages through which these biases emerge.

The present study aimed to determine when motor-related information becomes integrated into perceptual decision formation by examining the temporal dynamics of neural processing underlying the process of perceptual decision-making using EEG. Participants performed an orientation discrimination task in which two Gabor patches were presented simultaneously in the left and right hemifields, and they reported the Gabor perceived as more vertically oriented by pressing one of two pressure-sensitive response buttons with their dominant index finger. The motor effort required to report each choice was indicated by a force cue surrounding each Gabor patch, with the mapping between effort and spatial location blocked. Psychometric curve fitting was used to quantify motor effort-induced decision bias, and based on previous findings (Assarioti et al., 2025; Burk et al., 2014; Hagura et al., 2017; Marcos et al., 2015), we expected motor effort to bias perceptual decisions toward the low-effort option. Neurophysiologically, we measured: 1) parieto-occipital alpha power contralateral to low-effort relative to high-effort options to examine if spatial attention was biased toward the low-effort option; 2) P1 amplitudes and N1 amplitudes over the occipital cortex contralateral to low-effort relative to high-effort options, to examine if motor effort influenced early sensory processing; 3) CPP amplitude proceeding response execution for low-effort relative to high-effort choices to examine if motor effort influenced later-stage evidence accumulation. To further examine whether these neural effects predicted decision behavior, we looked at whether individual differences in neural modulation by motor effort correlated with participants’ decision bias, and at the single-trial level, whether neural activity favoring the low-effort option was associated with an increased likelihood of choosing that option.

## 2. Methods

### 2.1 Participants

Thirty-two healthy adults were recruited through the university’s online subject pool for participation in the study. All participants were right-handed, had normal or corrected-to-normal vision, and reported no history or current diagnosis of neurological, psychiatric, or musculoskeletal disorders. One participant was excluded due to an excessive number of rejected trials with insufficient fixation (based on concurrently collected eye-tracking data), and an additional seven participants were excluded based on visual inspection of their individual psychometric curves (see Section 2.5.1), resulting in a final sample of 24 participants (16 females; median age = 20 years, range = 18–26 years). All participants provided written informed consent prior to participation and received course credits as compensation. The study received ethics approval from the Scientific and Ethical Review Board of the Faculty of Behavioral and Movement Sciences at Vrije Universiteit Amsterdam (VCWE-2025-185).

### 2.2 Stimuli and materials

The experiment was programmed in MATLAB R2023b (The MathWorks, Natick, Massachusetts). Stimuli were generated and presented using the Psychophysics Toolbox Version 3.0.14 (Brainard, 1997; Pelli, 1997) on an ASUS ROG Strix XG248Q monitor (23.8″ diagonal, 1920 × 1080 resolution, 240 Hz refresh rate; physical dimensions: 526 mm × 296 mm), with the task setup illustrated in **Figure 1A**. Responses were collected using a pressure-sensitive USB response pad which registers input as applied force exerted against a rigid button (**Figure 1B**; The Black Box ToolKit Ltd., Sheffield, UK). Left and right responses were mapped to the first and second buttons, respectively. The applied force was continuously recorded, and responses were registered when applied force exceeded a predefined threshold (see below).

**Figure 1.**
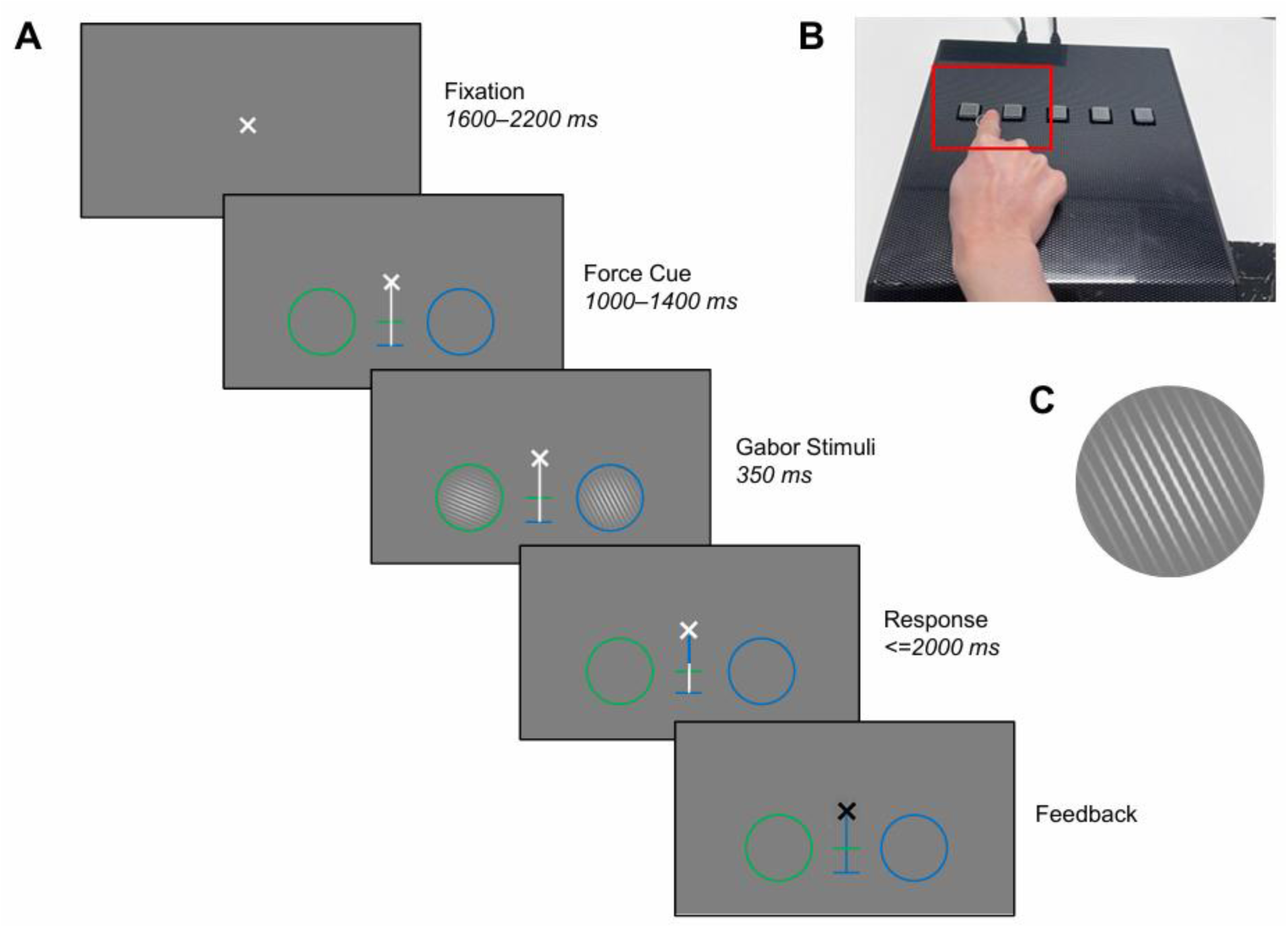
**(A)** Trial sequence. Each trial began with only a fixation cross presented centrally on the screen, followed by a force cue consisting of two colored circles (green = low effort; blue = high effort). Two Gabor patches were then presented centrally within the circles. Participants indicated which Gabor was more vertically oriented by pressing the corresponding button (see B). While pressing, visual feedback of the exerted force was provided via a gradual change in color of the vertical white bar matching the selected option downwards from the fixation cross until it reached the matching horizontal bar (in the illustrated example, a right button press resulted in blue filling toward the lower blue bar). The fixation cross turned black when the response was successfully registered and red if no response was registered within the response window. **(B)** Pressure-sensitive USB response pad used to register left and right responses through the first and second buttons, respectively. Participants rested the index finger of their dominant hand between the two buttons (marked by a sticker) and moved it to respond as required. **(C)** Example Gabor patch oriented at 20° anticlockwise from vertical, consisting of a sine-wave grating multiplied by a circular Gaussian envelope.

All stimuli were presented against a mid-grey background (RGB: 127, 127, 127). Among these was a white X-shaped fixation cross (RGB: 255, 255, 255), which was presented 1° of visual angle above the screen center, consisting of two diagonal lines (36 pixels in length, 8 pixels in thickness). A vertical white line (RGB: 255, 255, 255; 6 pixels in thickness) was presented at the screen midline spanning from 0.125° to 9° below fixation. Two horizontal colored lines (2.4° in width, 6 pixels in thickness) intersected this vertical line: the upper line was displayed in green (RGB: 0, 255, 0) at 4.5° below fixation, and the lower line was displayed in blue (RGB: 0, 0, 255) at 9° below fixation. Two circles (one green, one blue) were positioned 6° below and 5.3° left and right from the fixation cross, corresponding to an eccentricity of 8°. We will refer to the combination of colored circles and horizontal lines as the force cue (see section 2.3). Within each circle, a centrally positioned Gabor patch was briefly presented during each trial, scaled to 95% of the circle diameter. The Gabor patches (see example stimulus in **Figure 1C**) had a Michelson contrast of 0.8, a spatial frequency of approximately 2.8 cycles per degree, and a phase of 0°. They consisted of sine-wave gratings with 16 cycles across the patch multiplied by a circular Gaussian envelope (aspect ratio = 1, SD = 8 cycles). The orientation of the Gabor patches differed across trials.

### 2.3 Experimental procedure and design

The experiment was conducted in a quiet room with dim, constant artificial lighting, with the researcher located in an adjacent room. Participants were seated in a chair behind a table with a computer monitor at a viewing distance of approximately 70 cm. The chair height was adjusted to ensure physical comfort and optimal visibility of the display. As shown in **Figure 1B**, participants were instructed to rest the index finger of their dominant hand between the left and right buttons (marked by a sticker) while keeping their arms comfortably positioned on the table, and to respond by pressing one of the two buttons according to task requirements. The full experimental session lasted approximately 75 minutes, comprising approximately 30 minutes of preparation time (including task instructions and EEG setup) and 45 minutes of task execution (4 blocks, with short breaks provided between blocks to minimize fatigue).

Participants were instructed to keep their eyes on the fixation cross throughout the experiment and to make a decision on each trial regarding which of two Gabor patches was more vertically oriented by pressing the corresponding left or right button (see trial sequence details in **Figure 1A)**. Each response option required either low or high effort (force threshold), as indicated by the color of the circles surrounding the stimuli. This force cue was fixed: green = low effort; blue = high effort). The distance of the colored lines from the fixation cross (**Figure 1A**) thus corresponded to the effort required. The mapping between stimulus location and effort level was fixed within each block. There were two effort-mapping conditions: One with the low effort on the left and the high effort on the right (A) and one with the reverse mapping (B). Hereafter, these conditions are referred to as “low effort: left” and “low effort: right” respectively. At the beginning of each block, participants received instructions on the current mapping (e.g., “*During this block, the LEFT response requires LOW effort, and the RIGHT response requires HIGH effort*”). Each participant completed both mappings in an ABBA or BAAB sequence, counterbalanced across participants. Each block comprised 150 trials, resulting in a total of 600 trials.

Each trial began with the fixation cross presented alone on the screen for a random duration between 1600 and 2200 ms, followed by the force cue for a random duration between 1000 and 1400 ms. Two Gabor patches were subsequently simultaneously presented centrally within the circles for 350 ms. Participants responded as quickly and accurately as possible to which Gabor patch was more vertically oriented by pressing the corresponding button with the index finger of their dominant hand. Because the two response options required different levels of force, visual feedback was provided upon pressing the response key via a gradual downward color filling of the vertical white bar; the color matched the selected option and the downward extent indicated the exerted force. The color filled the vertical bar downwards from the fixation cross until the horizontal bar of the same color. The rate of filling was determined by two parameters: (1) the force threshold required to initiate filling and (2) the number of pixels traversed per unit of applied force (N). The low-effort (green) option required a threshold of 0.3 N and a gain of 2000 pixels per N, whereas the high-effort (blue) option required a threshold of 3 N and a gain of 200 pixels per N. This resulted in slower filling when choosing the high-effort option compared to the low-effort option. The response window was limited to 2000 ms. The fixation cross turned black when a response was successfully registered and turned red when no response was made within the response window (i.e. when no button was touched or when the applied force did not reach the threshold).

The two Gabors had a mean orientation of 45° anticlockwise for every trial. The orientation difference between the two Gabor patches was varied using a staircase procedure (Cornsweet, 1962). The orientation of each Gabor patch could vary at 1° resolution. The orientation difference was defined as the orientation of the left Gabor minus the orientation of the right Gabor. Within each block, two independent staircases were applied with one starting at an orientation difference of 40° and the other at −40°. The staircases were randomly interleaved and equally distributed across trials (75 trials per staircase). Each staircase followed a 2-up-2-down rule based on participants’ choices. For instance, considering an orientation difference of 40° (i.e. 25° and 65°), a right choice on that trial would lead to a decrease in the difference to 38° (26° and 64°) on the next trial of that staircase, whereas a left choice would lead to an increase to 42° (24° and 66°). In practice, the orientation difference in the experiments ranged between −40° and 44°.

Before they started the experiment, participants completed a practice session of 30 trials in one of the two effort-mapping conditions to familiarize themselves with the task. Feedback was provided after each response: “*correct*” in green for accurate responses; “*error*” in red for incorrect responses; “*too slow*” in white if no response was made within the 2000 ms response window; and “*press harder*” in white if a response was made but the applied force was insufficient to reach the threshold (i.e. for the color fill to reach the horizontal bar). This feedback was only presented in the practice session and not during the experiment. Participants were instructed to sit still and to keep their eyes on the central fixation cross throughout the practice.

### 2.4 EEG and eye-tracking acquisition

Electroencephalography (EEG) was recorded using a 64-channel BioSemi ActiveTwo system (BioSemi, Amsterdam, The Netherlands) at a sampling rate of 512 Hz. Electrodes were positioned according to the international 10–10 system, covering the entire scalp. During EEG acquisition, signals were referenced to the common mode sense (CMS) and driven right leg (DRL) electrodes, with scalp electrode-skin impedances maintained below 10 kΩ. Vertical (VEOG) and horizontal (HEOG) electrooculograms were recorded using electrodes placed approximately 2 cm above and below the right eye to monitor blinks and vertical eye movements, and at the outer canthi of both eyes to monitor horizontal saccades. Offline, EEG data were re-referenced to the average of all electrodes. Eye-tracking data were additionally collected at a 1000 Hz sampling rate using an EyeLink 1000 system (SR Research), calibrated at the beginning of the experiment and used to monitor participants’ fixation. Triggers were sent to both the EEG and eye-tracking devices to ensure synchronization with stimulus presentation.

### 2.5 Data Analyses

#### 2.5.1 Data cleaning and exclusion

All analyses were conducted using MATLAB R2023b unless specified otherwise. Data quality was assessed for each participant based on eye-tracking measures of fixation quality and task performance assessed by psychometric curve fits. Eye-tracking data were first converted to ASC format and then pre-processed using the open-source toolbox FieldTrip (version 2024-07-04; Oostenveld et al., 2011). The data were segmented on a trial-by-trial basis from force cue onset until feedback (i.e., when the fixation cross turned black or red). For each trial, a fixation center was determined using a baseline period defined as −300 to 0 ms relative to Gabor onset, by computing the median horizontal and vertical gaze coordinates across samples. This fixation center was then compared with each gaze sample in the full trial by computing the Euclidean distance to quantify fixation quality. Gaze samples falling within 2° of visual angle (approximately a radius of 90 pixels) were considered to reflect good fixation, and trials were excluded if fewer than 75% of samples fell within this range. Trials were also excluded if more than 50% of samples had no registered gaze points for example due to eye closure during the baseline period or across the entire trial. One participant was excluded due to an excessive proportion of rejected trials (∼30% of total trials). For the remaining participants the mean number of rejected trials was 47.7 (range = 0–158), equaling 7.9% (range: 0%–26.3%) of all trials. Trials rejected based on eye-tracking measures were excluded from all subsequent behavioral and EEG analyses.

Next, psychometric curves were estimated using *psignifit* version 4.0, a MATLAB toolbox for fitting parametric models to behavioral data, to quantify how stimulus intensity influences choice behavior (Fründ et al., 2011; Wichmann & Hill, 2001). Psychometric curves were fitted separately for each effort-mapping condition (low effort: left vs. low effort: right) for each participant, modelling the proportion of rightward choices as a function of the orientation difference between the two Gabor patches. Each function was modelled using a cumulative Gaussian distribution with all trials equally weighted. Four parameters were extracted from each fit: (1) the threshold, defined as the orientation difference at which 50% of choices were rightward; (2) the width, measuring the range of stimulus intensities required for the participant’s performance to change from the 5th to the 95th percentiles of the unscaled cumulative Gaussian; (3) the lapse rate, reflecting the proportion of stimulus-independent random errors made by the participant, corresponding to the symmetric upper and lower asymptotes of the function; and (4) the slope, indicating the rate of change in choice probability at the midpoint of the psychometric curve. We excluded participants with poor task engagement if any psychometric fit exhibited a lapse rate greater than 10%. Additionally, all fits were visually inspected to ensure that the fitted curves closely matched the observed data. These procedures resulted in the exclusion of seven participants (see **Figure 2** for examples of good and poor task performance).

**Figure 2.**
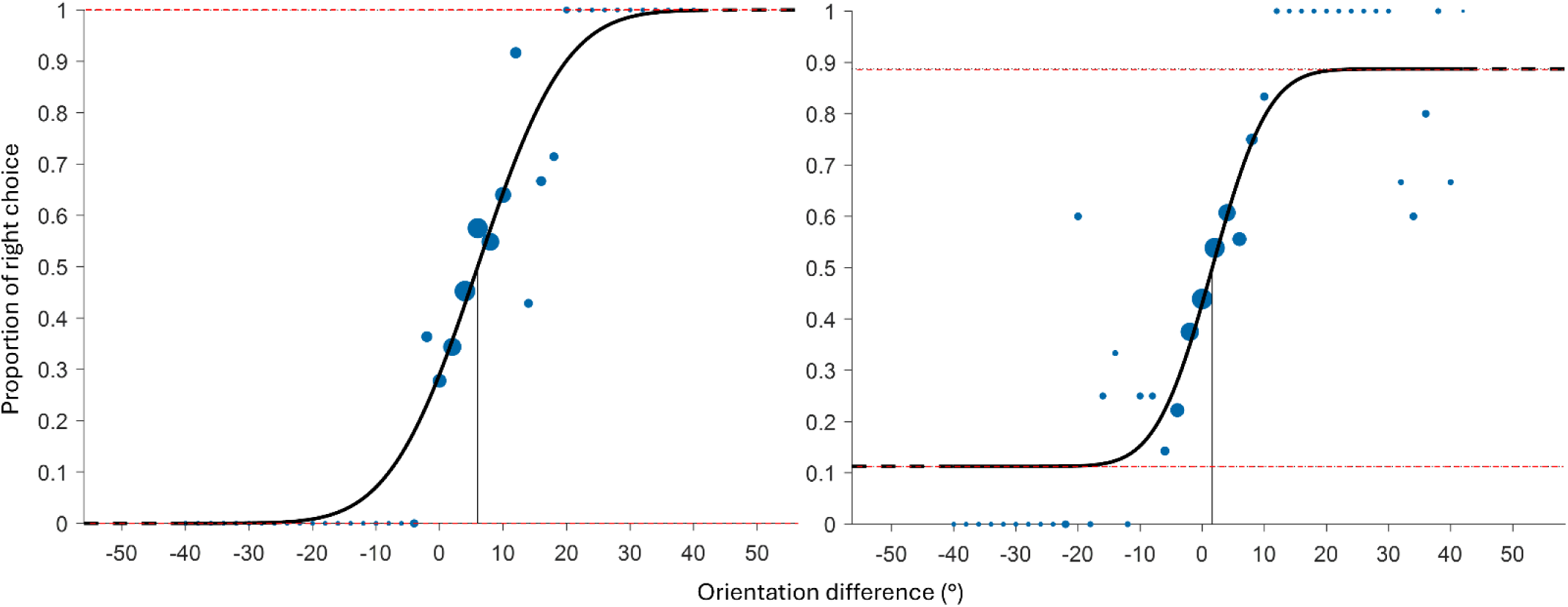
Example data (blue dots) and the best fitting psychometric functions (black curves) demonstrating good and poor decision-making task performance. The thin vertical black lines indicate the fitted threshold of each curve, corresponding to the orientation difference at which 50% of choices were rightward. Dashed red lines represent the fitted lapse rates, capturing deviations from perfect performance at the extremes of the stimulus range. Blue dots indicate the proportion of rightward choices at each orientation difference between the two Gabor stimuli, with dot size proportional to the square root of the number of trials at each level. The transition from solid to dashed in the black curves indicates the range of presented orientation differences. The left panel shows an example of good task performance characterized by a steep slope and low lapse rate, whereas the right panel shows an example of poor task performance with a high lapse rate (11.3%) and noticeable deviations of data points from the fitted curve. Participants showing poor fits comparable to this example were excluded.

#### 2.5.2 Behavioral analyses

Trials in which no response was registered were excluded, resulting in a mean of 1.8 rejected trials per participant (range: 1–24). To examine overall decision-making performance, reaction time (RT) was determined for each trial as the time relative to the onset of the Gabor presentation at which the continuously applied force for one of the two response options exceeded 0.01 N. Data were divided into four combinations of effort mapping (low effort: left vs. low effort: right) and choice (left vs. right). Median RT across trials was computed for each participant and each combination at each orientation difference and then averaged across all participants to obtain group-level RT estimates. To formally test RT differences across effort-mapping conditions and choices, trial-level RTs were analyzed as an exploratory analysis using a linear mixed-effects model with effort mapping, choice, the interaction between effort mapping and choice, and absolute orientation difference (a proxy for task difficulty) as fixed effects with a random intercept for participants.

To quantify decision bias due to motor effort, we compared the estimated threshold parameter (i.e., the orientation difference at which 50% of choices were rightward) of the psychometric curves fitted separately for the two effort-mapping conditions. For each fitted psychometric curve, a positive threshold reflects a bias towards leftward choices, whereas a negative threshold reflects a bias towards rightward choices. To obtain a single measure of decision bias independent of effort-mapping configuration, we subtracted the threshold in the low effort: right condition from the threshold in the low effort: left condition. A positive decision bias corresponds to a bias towards the low-effort option, indicating that participants required less evidence to choose the low-effort option. To test whether motor effort biased perceptual decisions toward low-effort options, decision bias was computed per participant and tested against zero using a Wilcoxon signed-rank test. As a supplementary analysis, drift-diffusion models were fitted to participants’ choice and RT data to examine the computational mechanisms underlying the motor effort–related decision bias. Full model specification and fitting procedures, as well as modeling results are reported in the Supplementary Methods.

#### 2.5.3 EEG analyses

##### 2.5.3.1 EEG preprocessing

All EEG analyses were performed using FieldTrip (version 2024-07-04; Oostenveld et al., 2011). Raw EEG data were first segmented into stimulus-locked epochs from −4 to 4.5 s relative to Gabor onset, which was sufficiently long to cover the full duration of each trial for subsequent re-segmentation. Vertical and horizontal EOG signals were constructed using bipolar derivations and were included together with the EEG data to facilitate visual artifact rejection. The epoched data were band-pass filtered between 0.1 and 70 Hz using a second-order bidirectional Butterworth filter, and line noise was removed using a notch filter (49–51 Hz). Visual artifact rejection was performed in two stages. First, prior to independent component analysis (ICA), trials containing large movement artifacts were removed, and noisy channels that could interfere with ICA decomposition were identified and interpolated using neighboring channels (mean = 0.5 channels per participant, range: 0–4). The data were then re-referenced to the average of all channels. ICA was conducted using the runica algorithm to identify and remove components reflecting non-neural activity, including ocular and muscle artifacts, aided by vertical and horizontal EOG signals. The ICA components were visually inspected, and components reflecting typical eye blinks, saccades, cardiac activity, and muscle artifacts were manually identified and removed. On average, 16.9 components were removed per participant (range: 7–27). To maintain consistency with the behavioral analyses (see Section 2.5.2), trials without a registered response were excluded. Given that the primary research question concerned neural activity spanning stimulus processing through response commitment, the data were re-segmented into a stimulus-locked epoch (−2 to 2.75 s relative to Gabor onset) and a response-locked epoch (−2.5 to 2 s relative to response onset). Finally, a second round of visual artifact rejection was performed to discard trials with any residual artifacts.

##### 2.5.3.2 Time–frequency analysis

Time–frequency (TF) analysis was conducted on the stimulus-locked epoch to examine pre-stimulus (during force cue presentation) parieto-occipital alpha activity, which indexes attentional allocation and processing. Analyses focused on parieto-occipital electrodes grouped by hemisphere (left: PO7, O1, PO3; right: PO8, O2, PO4), consistent with prior attention research (e.g., Duncan et al., 2025; Thut et al., 2006; Wang et al., 2019). TF representations were computed using a multi-taper approach (5–40 Hz, 1 Hz steps) with 5-cycle adaptive windows, 0.4 × frequency smoothing, and a time window from −1.5 to 2.5 s (50 ms steps). TF estimates were computed separately for each hemisphere, effort-mapping condition, and participant, and were normalized using a decibel (dB) log-ratio transformation to the −500 to −100 ms time interval relative to force cue onset. For each participant, the TF representation contralateral to the low-effort option was computed by averaging TF estimates across trials and selected electrodes for the left hemisphere for trials in which the low-effort option appeared in the right visual field and across trials and selected electrodes for the right hemisphere for trials in which the low-effort option appeared in the left visual field; the same procedure was applied for the high-effort option. To statistically test whether motor effort influences attentional allocation prior to evidence accumulation, paired-samples t-tests were conducted to compare averaged alpha power within the 8–12 Hz frequency range and the −1000 to 0 ms window relative to stimulus onset, selected based on the presentation duration of the force-cue, contralateral to low- versus high-effort options.

##### 2.5.3.3 Event-related potential analysis

Analyses of event-related potentials (ERPs) were conducted to characterize neural signatures of early sensory encoding, specifically the P1 and N1 components (Hillyard & Anllo-Vento, 1998), as well as the CPP associated with sensory evidence accumulation during decision formation (Kelly & O’Connell, 2013; O’Connell et al., 2012). For the P1 and N1 components, time-locked analyses were performed on the stimulus-locked epoch separately for each effort-mapping condition and quantified at parieto-occipital electrodes grouped by hemisphere (left: PO7, O1, PO3; right: PO8, O2, PO4), in line with prior visual ERP research (e.g., Di Russo et al., 2002; Hillyard & Anllo-Vento, 1998). Data were baseline-corrected on each trial by subtracting the mean voltage in the −200 to 0 ms pre-stimulus interval. Mean ERP amplitudes were calculated separately for each hemisphere, participant, and effort-mapping condition, with contralateral ERPs to the low-effort option computed by averaging ERPs across trials and selected electrodes for the left hemisphere for trials in which the low-effort option appeared in the right visual field and across trials and selected electrodes for the right hemisphere for trials in which the low-effort option appeared in the left visual field; the same procedure was applied to compute contralateral ERPs for the high-effort option. To test whether motor effort influences early visual processing of decision options, paired-samples t-tests were conducted to assess the amplitude difference in P1 and N1 contralateral to low- versus high-effort options averaged within established time windows (80–130 ms for P1; 150–200 ms for N1; e.g., Di Russo et al., 2002; Hillyard & Anllo-Vento, 1998).

The CPP component was constructed using the response-locked epoch separately for trials in which low-effort and high-effort options were chosen and averaged across central midline electrodes (FCz, Cz, CPz), where this signature is typically most pronounced and commonly examined in prior decision-making research (e.g., Kelly & O’Connell, 2013; O’Connell et al., 2012). Data were baseline-corrected on each trial using the −200 to 0 ms pre-stimulus interval. Mean ERP amplitudes were computed for each participant and subsequently grand-averaged across participants for visualization. Based on inspection of the CPP waveform (see **Figure 7A**), which showed a plateau beginning approximately 200 ms prior to response onset, and to test whether motor effort influences late-stage evidence accumulation, paired-samples t-tests were conducted to compare average CPP amplitude between trials with low- and high-effort choices within the −200 to −50 ms time window relative to response onset.

#### 2.5.4 Brain-behavior relationships

To gain insight into the stage at which motor effort-related decision bias emerges during decision formation, we tested whether the computed neural signatures were associated with decision behavior. To this end, Spearman correlations were conducted at the between-subject level and logistic mixed-effects analyses were conducted at the within-subject (single-trial) level. At the between-subject level, Spearman correlations were conducted between decision bias, quantified as the psychometric threshold shift between the two effort-mapping conditions, and (a) the difference in normalized alpha power over parieto-occipital cortex contralateral to low- versus high-effort options, (b) the difference in P1 and N1 amplitudes over parieto-occipital cortex contralateral to low- versus high-effort options, and (c) the difference in CPP amplitude between low- and high-effort choices. At the within-subject level, logistic mixed-effects models were performed on trial-wise choice (low-effort choice: 0/1). Fixed effects included (a) z-scored trial-level differences in alpha power contralateral to low- versus high-effort options, (b) z-scored trial-level differences in P1 and N1 amplitudes contralateral to low- versus high-effort options, and (c) z-scored trial-level CPP amplitude. Participants were included as random intercepts for each neural predictor.

## 3. Results

### 3.1 Motor effort biases perceptual decisions toward low-effort options

Participants’ decision-making performance was first examined by analyzing RTs as a function of effort mapping (low effort: left vs. low effort: right), choice (left vs. right) and orientation difference. As shown in **Figure 3A**, RTs increased as orientation differences approached zero, indicating slower responses for more difficult stimuli. Choices followed the expected pattern: left choices predominated when orientation differences were negative (i.e., the left Gabor was more vertical), whereas right choices predominated when orientation differences were positive (i.e., the right Gabor was more vertical). Interestingly, RTs were also slower for large orientation differences, consistent with observations reported in Assarioti et al. (2025), suggesting an initial adjustment period at the beginning of each block. As an exploratory analysis, trial-level RTs were analyzed using a linear mixed-effects model with choice, effort mapping, their interaction, and absolute orientation difference as fixed effects, and random intercepts for participants. RTs decreased with increasing absolute orientation difference (β = −0.0037, SE = 0.0002), indicating faster responses for easier stimuli. There was no main effect of effort mapping (β = −0.004 ± 0.005) but a main effect of choice (β = −0.012, SE = 0.005). There was an interaction between choice and effort mapping (β = 0.041, SE = 0.007), indicating that RTs depended on the alignment between choice side and effort mapping. Specifically, when the low-effort option was on the left, left choices were slower (median = 0.75 s, IQR = [0.61, 0.93] s) than right choices (median = 0.73 s, IQR = [0.59, 0.90] s); when the low-effort option was on the right, right choices were slower (median = 0.78 s, IQR = [0.63, 0.95] s) than left choices (median = 0.75 s, IQR = [0.61, 0.92] s). One possible explanation is that participants executed responses toward the high-effort options more vigorously, leading to earlier crossing of the response threshold and thus earlier detection of RT.

**Figure 3.**
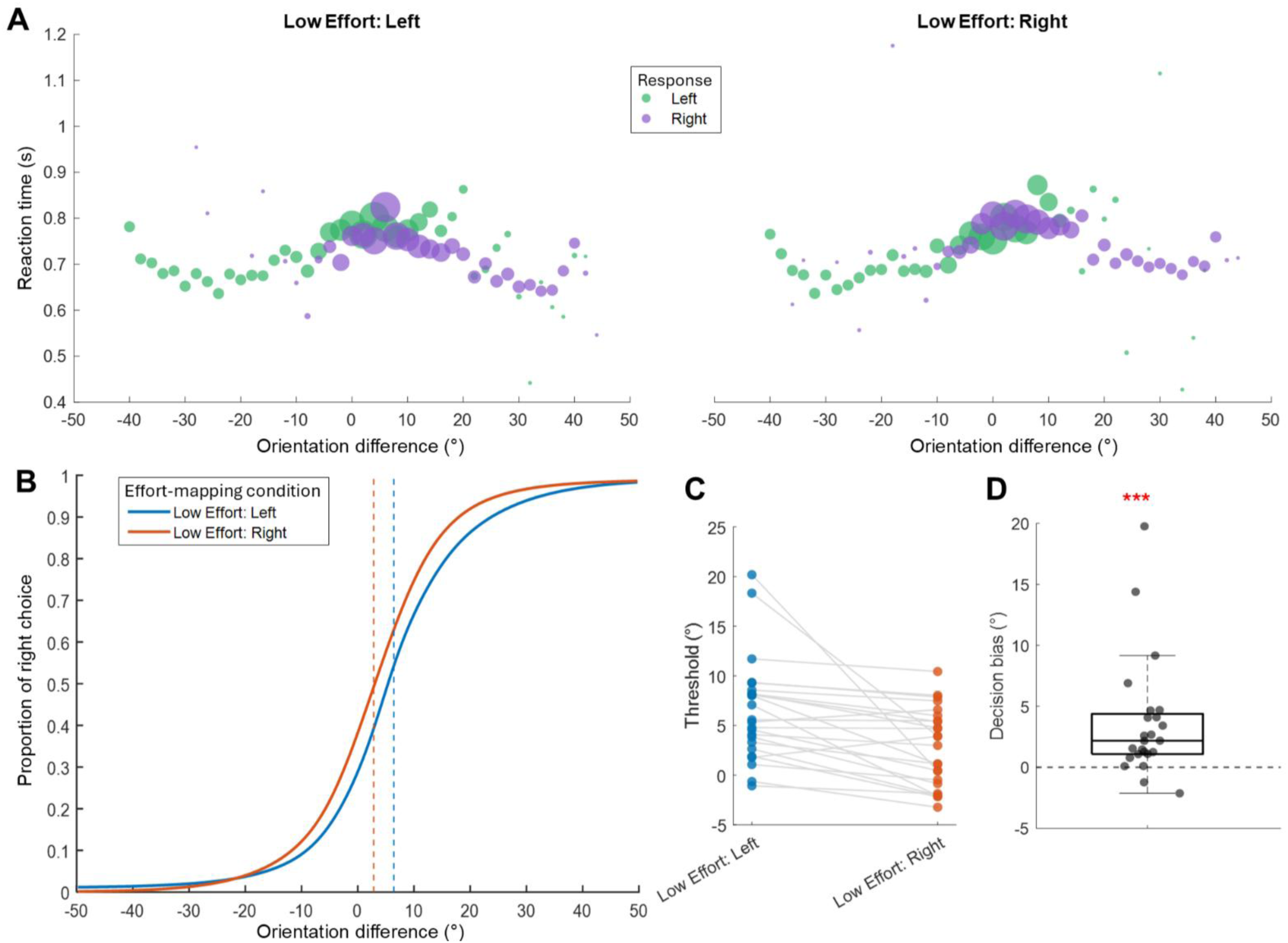
**(A)** Group-averaged median RTs plotted against orientation difference across effort-mapping conditions and choice. Orientation difference was defined as the orientation of the left Gabor minus that of the right Gabor, with positive values indicating that the right Gabor was more vertically oriented and negative values indicating that the left Gabor was more vertically oriented. Task difficulty increased as the orientation difference approached zero. The area of each data point represents the proportion of trials associated with each orientation difference and combination of effort mapping and choice. **(B)** Group-averaged psychometric curves fitted for each effort-mapping condition, showing the proportion of rightward choices as a function of orientation difference. Dashed vertical lines indicate threshold values at which the probability of a rightward choice is 50%. Larger thresholds indicate that more sensory evidence was required to select the right choice option. **(C)** Thresholds for each effort-mapping condition plotted per participant. Higher thresholds when the low-effort option was on the left compared to when it was on the right indicate that more evidence was required to reach 50% rightward choices in this effort-mapping condition. **(D)** Decision bias, quantified as the difference in thresholds between effort-mapping conditions, plotted for individual participants. Positive values indicate a bias toward low-effort choices, whereas negative values indicate a bias toward high-effort choices. The line in the center of the boxplot represents the median, the box indicates the interquartile range, and the whiskers indicate the most extreme data points within 1.5 times of the interquartile range below Q1 and above Q3. The three asterisks indicate a significant difference (*p* <.001).

To test whether motor effort biases perceptual decisions, psychometric curves were fitted separately for each effort-mapping condition, and thresholds were computed to represent the choice proportion at 50% rightward choices. All other parameter estimates from the psychometric functions for each effort-mapping condition, averaged across participants, are reported in Supplementary Table S1. As shown in the group-averaged curves (**Figure 3B**), thresholds were larger when the low-effort option appeared on the left (Mean = 6.4°, SE = 1.1°) than when it appeared on the right (Mean = 2.8°, SE = 0.8°), indicating that more sensory evidence was required to reach 50% rightward choices when rightward choices were associated with higher effort. This effect was consistent at the individual level (**Figure 3C**). Interestingly, we observed a general leftward choice bias, indicated by the positive threshold shift in both effort-mapping conditions, consistent with Assarioti et al. (2025). This bias may be related to the Gabor patches always presented anticlockwise. To provide a single measure independent of effort mapping, decision bias was quantified as the difference in thresholds between effort-mapping conditions, and was found to be significantly greater than zero (Wilcoxon signed-rank test: median difference = 2.18°, W = 283, z = 3.79, *p* < .001; **Figure 3D**), indicating that motor effort biased perceptual decisions toward low-effort options.

### 3.2 Pre-stimulus alpha power reflects low-effort attentional bias but does not predict decision bias

To examine whether motor effort modulates preparatory attentional allocation prior to decision stimulus presentation, we analyzed pre-stimulus alpha power over parieto-occipital regions (left: PO7, O1, PO3; right: PO8, O2, PO4) contralateral to low- and high-effort options. TF representations were computed by averaging activity from electrodes contralateral to the corresponding option for each participant and baseline-normalized using the −500 to −100 ms time interval relative to force cue onset. As shown in **Figure 4A**, grand-averaged normalized alpha power time series (8–12 Hz) in parieto-occipital cortex contralateral to the low-effort option showed a greater decrease compared to contralateral to the high-effort option prior to stimulus onset and extending into early stimulus processing. Mean baseline-normalized alpha power within the pre-stimulus interval (−1000 to 0 ms) was computed separately for parieto-occipital electrodes contralateral to low- and high-effort options for each participant (**Figure 4B**), and alpha power difference scores between these two options were subsequently calculated for each participant (**Figure 4C**). A paired-samples t-test revealed significantly lower alpha power contralateral to the low-effort option than contralateral to the high-effort option (t (23) = −3.05, *p* = .006), suggesting enhanced attentional allocation toward low-effort options prior to the presentation of the decision stimulus. For visualization, **Figure 4D** shows grand-averaged TF representations of the contrast between activity contralateral to low- versus high-effort options over parieto-occipital cortex, together with the scalp topography averaged across the alpha band (8–12 Hz) and the −1000 to 0 ms interval, revealing a clear decrease in alpha power localized over parieto-occipital regions.

**Figure 4.**
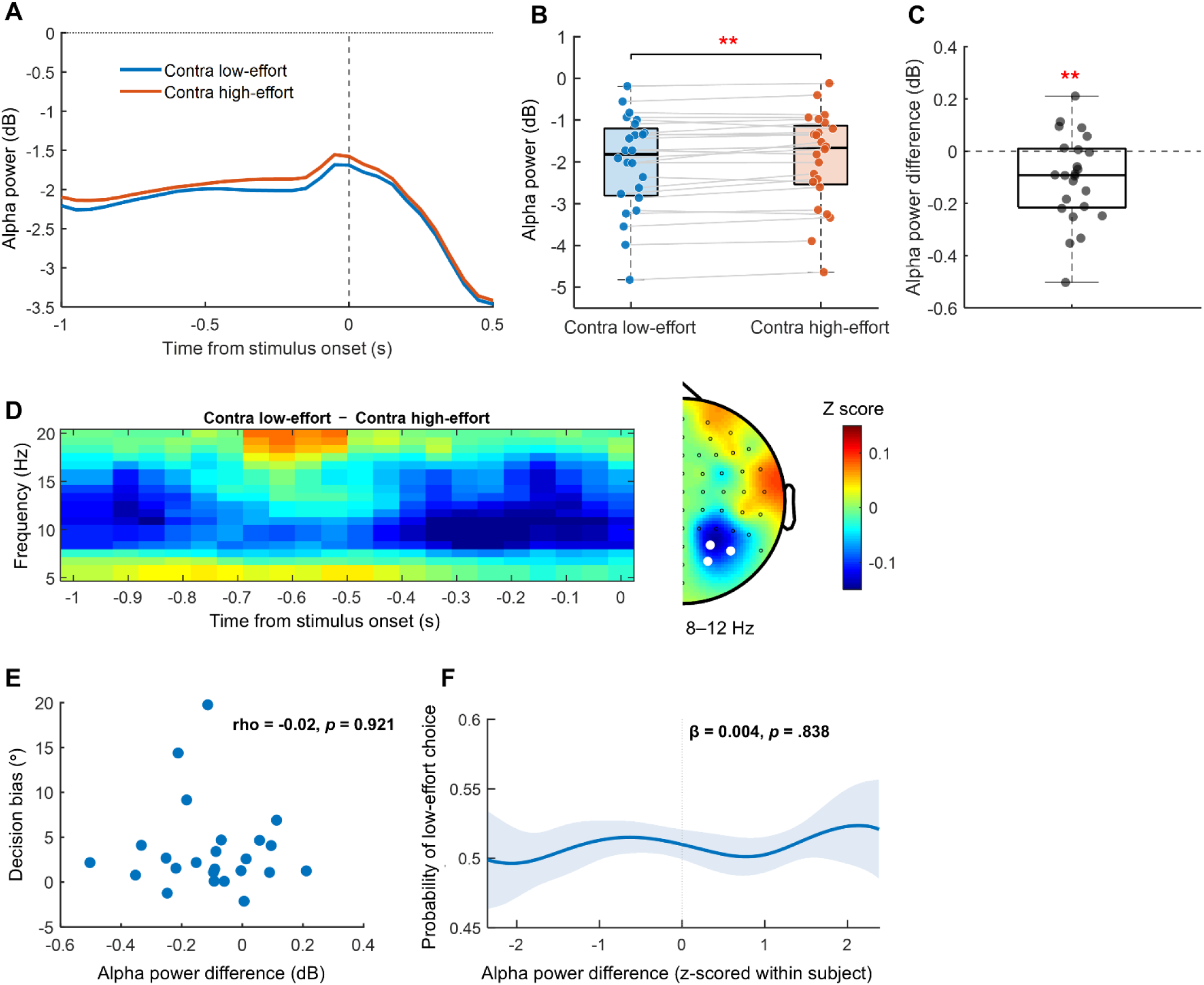
**(A)** Grand-averaged alpha power (8–12 Hz) time series over parieto-occipital cortex (left: PO7, O1, PO3; right: PO8, O2, PO4) computed contralateral to the low- and high-effort options, time-locked to stimulus onset and baseline-normalized to the −500 to −100 ms interval prior to force cue onset. **(B)** Normalized alpha power averaged within the −1000 to 0 ms pre-stimulus interval for each participant, shown separately for contra low- and high-effort options. The two asterisks indicate a significant difference (*p* <.01). **(C)** Baseline-normalized alpha power difference scores computed by subtracting alpha power contralateral to high-effort options from alpha power contralateral to low-effort options. Each black dot represents the difference between the corresponding blue and red dots for an individual participant in panel B. **(D)** Grand-averaged TF representations (left) of the contrast between parieto-occipital activity contralateral to low- versus high-effort options, together with the scalp topography averaged across the alpha band (8–12 Hz) and the −1 to 0 s interval relative to decision stimulus onset (right). White dots indicate the parieto-occipital regions of interest. The color scale represents z-scores, with warmer colors indicating more positive values and cooler colors indicating more negative values. **(E)** Between-subject Spearman correlation between decision bias and the alpha power difference between contralateral low- and high-effort options. Each blue dot represents an individual participant. **(F)** Trial-level relationship between the probability of choosing the low-effort option and z-scored alpha power difference. For visualization, alpha-power values across the 1st–99th percentile range were plotted, and 200 equally spaced values were chosen within this range as x-axis. A Gaussian-weighted smoothing method (Van Leeuwen et al., 2019) was subsequently used. For each participant, trials with an alpha-power value closer to a given x-axis value were weighted more strongly in estimating choice probability, using a bandwidth of 0.4 z-score units. Smoothed curves were computed separately for each participant and then averaged across participants. Blue shaded regions indicate 95% confidence intervals across participants.

We further tested whether this alpha power difference (**Figure 4C**) predicts behavioral decision bias. At the between-subject level, a Spearman correlation revealed no significant association between decision bias quantified as the psychometric threshold shift between effort-mapping conditions and the difference in normalized alpha power contralateral to low- and high-effort options (rho = −0.02, *p* = .921; see **Figure 4E**). In addition, an absence of within-subject effects was observed at the trial level, as revealed by a logistic mixed-effects model predicting trial-wise choice (low-effort choice: 0/1) from the trial-level z-scored alpha power difference (β = 0.004, SE = 0.018, *p* = .838), with participants included as random intercepts (see **Figure 4F)**. These results suggest that although attention was more strongly allocated to the low-effort option prior to the presentation of the decision stimulus, this attentional bias did not explain the decision bias.

### 3.3 No evidence that effort modulates early sensory encoding (P1/N1)

To examine whether motor effort influences early visual encoding during perceptual decisions, we analyzed P1 and N1 components over parieto-occipital regions (left: PO7, O1, PO3; right: PO8, O2, PO4), contralateral to low- and high-effort options. ERPs were computed by averaging activity from electrodes contralateral to the corresponding option for each participant, with baseline correction applied on each trial using the −200 to 0 ms interval relative to stimulus onset. As shown in **Figure 5A**, grand-averaged ERPs in parieto-occipital cortex contralateral to low- and high-effort conditions largely overlapped, indicating no reliable difference between conditions. This was confirmed by paired-samples t-tests, which revealed no significant differences for either component (P1: t (23)= 0.85, *p* = .404; N1: t (23) = 1.13, *p*= .272). Scalp topographies were computed using amplitudes averaged within predefined time windows for P1 (80–130 ms) and N1 (150–200 ms) and across all trials, showing the expected positive deflection over occipital cortex for P1 and negative deflection for N1. This provided no evidence that motor effort modulated early sensory encoding of the decision stimulus.

**Figure 5.**
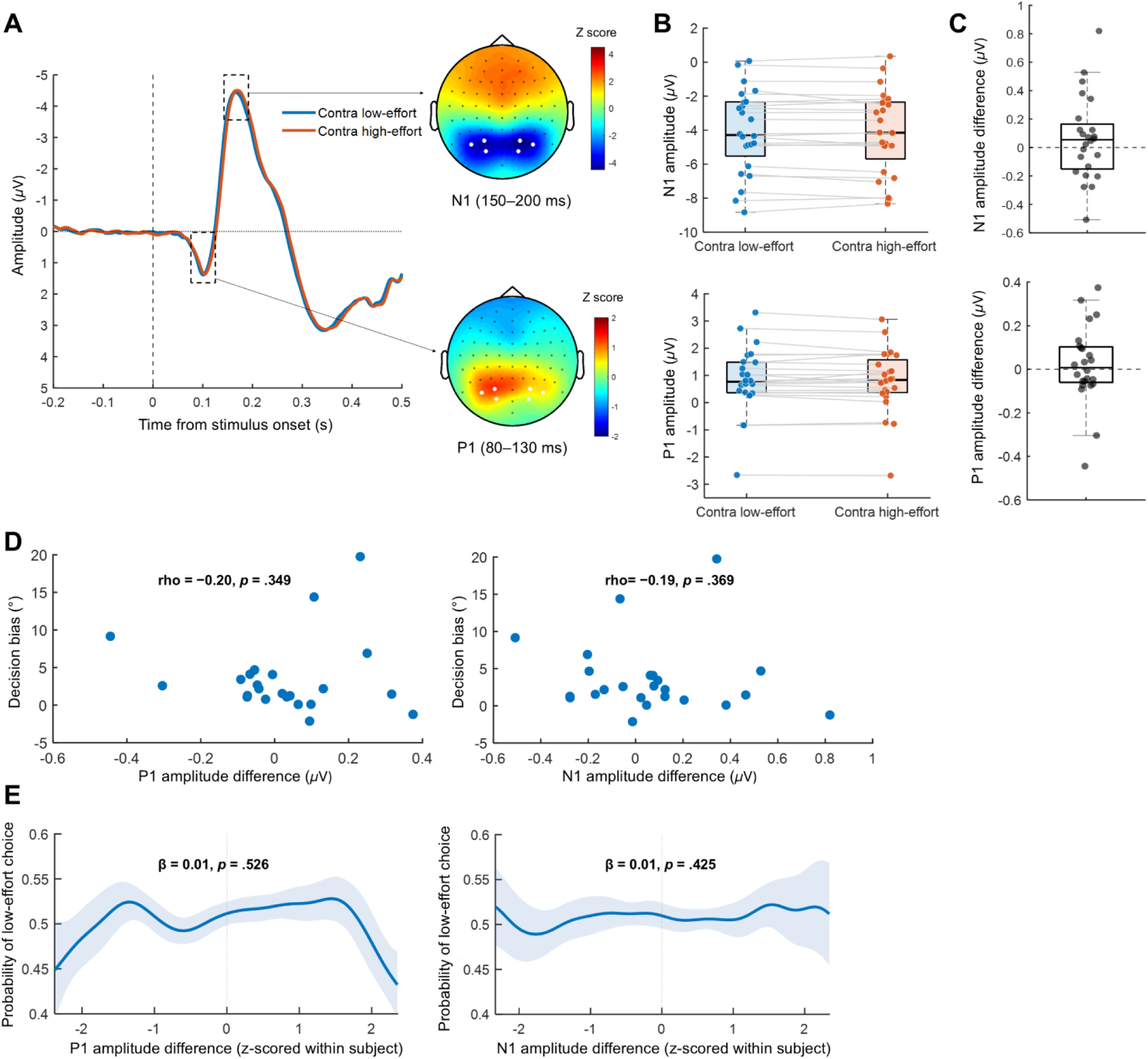
**(A)** Grand-averaged ERP time series averaged over parieto-occipital electrodes (left: PO7, O1, PO3; right: PO8, O2, PO4) contralateral to the low- and high-effort options, time-locked to stimulus onset and baseline-corrected to the −200 to 0 ms pre-stimulus interval. Dashed rectangles indicate the P1 and N1 components, with scalp topographies computed by averaging activity within predefined time windows for P1 (80–130 ms) and N1 (150–200 ms) collapsed across all trials. The color scale represents z-scores with warmer colors indicating more positive values and cooler colors indicating more negative values. **(B)** Mean amplitudes for the P1 and N1 components averaged within the predefined windows for each participant, shown separately for contra low- and high-effort options. **(C)** Mean amplitude difference for the P1 and N1 components computed by subtracting activity contralateral to high-effort options from activity contralateral to low-effort options. Each black dot represents the difference between the corresponding blue and red dots for an individual participant in panel B. **(D)** Between-subject correlations between decision bias and the P1 and N1 amplitude differences between parieto-occipital activity contralateral to low- versus high-effort options. Each blue dot represents an individual participant. **(E)** Trial-level relationship between the probability of choosing the low-effort option and z-scored P1/N1 amplitude difference. For visualization, Gaussian-weighted smoothing was applied using the same procedure as described in Figure 4F.

Despite the absence of motor effort modulation at the group level, we further tested whether differences in P1 and N1 amplitudes between parieto-occipital cortex contralateral to the low- vs. high-effort options explained inter- and intraindividual differences in the observed decision bias. Neither the between-subject Spearman correlations (P1: rho = −0.20, *p* = .349; N1: rho= −0.19, *p* = .369) nor the within-subject logistic mixed-effects models (P1: β = 0.01, SE = 0.02, *p* = .526; N1: β = 0.01, SE = 0.02, *p* = .425) revealed significant effects, suggesting no evidence that early visual processing contributed to motor effort–related decision behavior.

### 3.4 CPP exhibits greater amplitude prior to response execution for low-effort choices and predicts trial-level choice behavior

To examine whether motor effort modulates late-stage evidence accumulation during perceptual decisions, response-locked analyses were conducted to construct the CPP from central midline electrodes (FCz, Cz, CPz) for trials in which low-effort options were chosen and those in which high-effort options were chosen, with baseline correction applied on each trial using the −200 to 0 ms interval relative to stimulus onset. As illustrated in **Figure 6A**, the grand-averaged ERP time series showed an earlier and overall greater CPP build-up prior to response onset for trials in which the low-effort option was chosen compared to trials in which the high-effort option was chosen. Around 200 ms before the response, both choice types reached a plateau, indicating that evidence accumulation had stabilized. The scalp topography of the contrast (low- vs. high-effort choices) averaged within the −200 to −50 ms interval relative to response onset, demonstrated a clear central positivity. Mean CPP amplitude within this interval for trials with low-effort choices was found to be significantly higher than for trials with high-effort choices (t (23)= 3.24, *p* < .001, **Figure 6B**). This suggests that motor effort modulates late-stage sensory evidence accumulation such that the decision process accumulated more strongly toward choosing the low-effort option.

**Figure 6.**
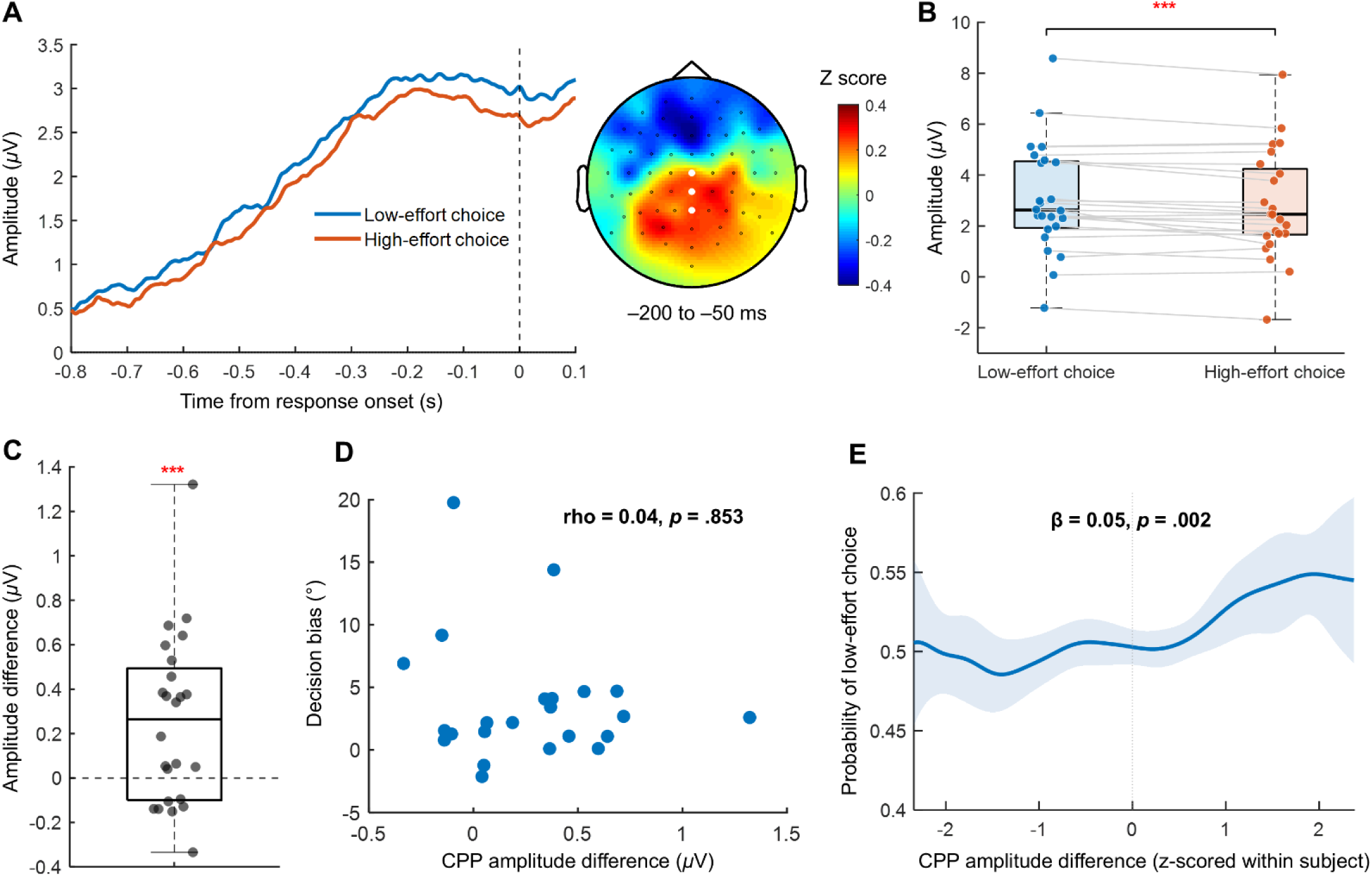
**(A)** Grand-averaged CPP time series (left) averaged over central midline electrodes (FCz, Cz, CPz) for trials in which the low- and high-effort options were chosen, time-locked to response onset and baseline-corrected to the −200 to 0 ms pre-stimulus interval. The scalp topography (right) shows the mean CPP amplitude difference between low- vs. high-effort choices, averaged over the −200 to −50 ms pre-response interval. The color scale represents z-scores, with warmer colors indicating more positive values and cooler colors indicating more negative values. **(B)** Mean CPP amplitudes within the −200 to −50 ms pre-response interval for low- and high-effort choices. The three asterisks indicate a significant difference (*p* < .001). **(C)** Mean CPP amplitude difference scores computed by subtracting CPP amplitudes on trials with high-effort choices from those on trials with low-effort choices. Each black dot represents the difference between the corresponding blue and red dots for an individual participant in panel B. **(D)** Between-subject correlations between decision bias and the CPP amplitude differences between low- versus high-effort options. Each blue dot represents an individual participant. **(F)** Trial-level relationship between the probability of choosing the low-effort option and z-scored CPP amplitude difference. For visualization, Gaussian-weighted smoothing was applied using the same procedure as described in Figure 5F.

We further examined whether motor effort–related differences in evidence accumulation accounted for the observed behavioral decision bias. A between-subject Spearman correlation tested whether the decision bias, quantified as the psychometric threshold shift between effort-mapping conditions, was associated with the CPP amplitude difference for −200 to 0 ms interval between low- and high-effort choices. No significant correlation was observed (rho = 0.04, *p* = .853; see **Figure 6D**). At the within-subject level, a logistic mixed-effects model predicting trial-wise choice (low-effort choice: 0/1) was conducted, with trial-level CPP amplitude (z-scored within participants) included as fixed effects and participants as random intercepts. This revealed a significant positive relationship between z-scored CPP amplitude and probability of low-effort choice (β = 0.05, SE = 0.02, *p* = .002), indicating that trials with higher CPP amplitude relative to participants’ own average value were associated with a higher likelihood of low-effort choices. These findings suggest that trial-wise CPP amplitude was associated with choice behavior and reflected a decision bias towards the low-effort option.

## 4. Discussion

The present study aimed to investigate when motor effort becomes integrated into perceptual decision formation by examining the temporal evolution of neural activity measured with EEG. We conducted an orientation discrimination task where participants were instructed to report the more vertically oriented Gabor patch among the two alternatives that appeared simultaneously in the left and right visual fields, by pressing one of the two corresponding pressure-sensitive response buttons. Motor effort required for reporting each choice was varied across blocks and indicated by force cues surrounding each Gabor patch. We found that perceptual decisions were biased toward the low-effort option. Motor effort modulated both pre-stimulus alpha-band power and CPP amplitude proceeding response execution, whereas no modulation was observed during early sensory encoding as indexed by the P1 and N1 components. Importantly, although motor effort influenced pre-stimulus alpha power, this effect was not associated with behavioral decision bias. By contrast, the CPP exhibited greater amplitude prior to response execution for low-effort choices and trial-level CPP amplitude significantly led to an increased likelihood of selecting the low-effort option. Together, these findings suggest that the motor effort-induced decision bias emerges during later-stage evidence accumulation rather than early sensory-attentional processing.

Our finding that perceptual decisions can be biased by motor effort is consistent with previous behavioral studies that varied reaching distance (Assarioti et al., 2025), movement angle (Marcos et al., 2015) or the physical resistance associated with each response (Hagura et al., 2017), all of which showed a consistent bias toward less effortful choices. It is also in line with findings that higher motor effort reduces the frequency of changes of mind (Burk et al., 2014). Our neural results are consistent with, but also extend those of Hagura et al. (2017) who used a similar psychometric function approach in a random-dot motion discrimination task in which exerted effort associated with each response was implicitly manipulated. They observed a decision bias toward the low-effort option, reflected by a shift in the psychometric threshold but not in the slope, indicating that motor costs biased perceptual decisions without altering sensory discrimination. They proposed that motor costs did not alter how sensory input is extracted and encoded in the visual system. Similarly, the psychometric threshold differed between effort-mapping conditions in the present study, and we further provided direct neural evidence by showing that early sensory processing, as indexed by the P1 and N1 components over occipital cortex, was not modulated by motor effort.

Despite the finding that motor effort did not appear to alter the quality of incoming sensory representations, our results showed that it modulated the preparatory state of the visual system, as reflected by reduced pre-stimulus alpha activity contralateral to the low-effort option. This is consistent with previous work suggesting that spatial attention within near space is action-oriented (Makin et al., 2010), as well as the active-inference framework suggesting that perception and attention are organized around how actions are predicted and controlled (Friston et al., 2017; Lageman et al., 2026). It may also be explained by the fact that effort mapping in our study remained constant within each block, allowing participants to be fully aware in advance which response would require more or less effort before stimulus onset. However, this preparatory attentional modulation was not associated with behavioral decision bias at either the between-subject or trial level, suggesting that although motor effort may bias attentional tuning toward the low-effort option before sensory evidence is presented, this preparatory bias is unlikely to be the primary mechanism underlying the observed behavioral bias.

Instead, the greater CPP amplitude preceding response execution for low-effort choices, together with the positive trial-level association between CPP amplitude and the probability of selecting the low-effort option, suggests that anticipated motor effort biased the evidence-accumulation process towards the low-effort option. In other words, motor effort does not alter what we perceive, but rather alters how perceptual information is processed during decision formation. This is consistent with the idea that decision making and action planning are considered closely coupled and, in some cases, evolve in parallel (Cisek, 2007; Cisek & Kalaska, 2010). Hence, motor information may be more likely to influence the decision process when individuals weigh the two options and map them onto potential action plans, rather than during the initial visual processing of how the options are presented. This finding is in line with earlier neurophysiological studies demonstrating that motor cortex activity might be actively involved in encoding decision variables instead of merely serving as a substrate for motor output (Cui et al., 2026; Klein-Flügge et al., 2016; Klein-Flügge & Bestmann, 2012; Michelet et al., 2010; Pape & Siegel, 2016; Urai & Donner, 2022), and further supports the framework of embodied perceptual decision-making (Cisek, 2007; Lepora & Pezzulo, 2015; Pastor-Bernier & Cisek, 2011). One point to note, however, is that if motor effort facilitates evidence accumulation toward the low-effort option, we should expect faster RT for low-effort choices. This was not, however, observed in the present study where RT was found to be faster for high-effort choices. A similar absence of an RT advantage for low-effort responses has also been reported in Marcos et al. (2015). This may be explained by the fact that RT in the present study was defined as the time at which the force threshold was crossed. Participants may have pressed the button more vigorously when reporting high-effort choices, leading to more rapid threshold crossing despite slower decision formation.

Despite the within-participant trial-level relationship between low-effort decision probability and CPP amplitude, the lack of a between-subject correlation between individual decision bias and mean CPP amplitude difference between low-and high-effort choices suggests that the way motor effort shapes evidence accumulation is a more momentary state-dependent phenomenon rather than one tied to stable individual traits. This is consistent with previous studies in which CPP is typically examined at the trial level and has been found to be a dynamic signal that predicts trial-level behavior in decision-making (Parés-Pujolràs et al., 2025; Van Vugt et al., 2019).

From a computational perspective, motor effort could influence perceptual decision formation in two ways: by shifting the starting point, such that evidence accumulation starts closer to one decision bound, or by altering the drift rate, such that evidence is accumulated more strongly over the time course of decision-making. Our DDM results (see details in Supplementary Results) showed that motor effort appeared to alter both starting-point and drift-rate parameters, and individual differences in motor effort-induced decision bias, quantified by the psychometric threshold difference between effort-mapping conditions, were associated with both motor effort-related starting-point and drift-rate parameters. There was, however, a potential trade-off between the two parameters, as suggested by their negative correlation, making it difficult to attribute the observed decision bias to either computational mechanism. These results are consistent with Hanks et al. (2011) who showed that prior probability can be integrated dynamically during evidence accumulation when the reliability of sensory evidence varies from trial to trial, rather than acting merely as an initial decision bias. However, they differ from several prior studies showing that decision bias was best explained by a static shift in the starting point of evidence accumulation, including work on motor costs (Hagura et al., 2017), prior probability and reward (Mulder et al., 2012) and neural perturbations (Ding & Gold, 2012; Hanks et al., 2006). One possible explanation for this discrepancy lies in the different modeling procedures. We fitted a model in which both motor effort-related drift-rate and starting-point parameters varied to individual participants’ data, which captured individual behavior, whereas previous studies fitted separate models to pooled data and conducted model comparisons (Hagura et al., 2017). Moreover, differences in task design may have also played a role: when task difficulty is fixed, decisions may rely more heavily on an initial bias, whereas when task difficulty varies across trials (as in the present study), greater weighting may be placed on evidence accumulated over the course of decision-making (Mulder et al., 2012).

Several limitations of the current study should be noted. First, because the force mapping in our study remained constant within each block, it is unclear whether similar effects would be observed if force mapping varied unpredictably across trials as in Assarioti et al. (2025). Knowing which side was more effortful throughout a block might have enabled participants to learn the motor-effort contingencies over time and get adapted to them, whereas trial-wise force mapping would require more rapid updating of action plans. Second, the motor effort manipulation was relatively explicit, as reflected by the large force difference between two options and explicit force cues. Future studies should examine whether similar decision biases arise under more implicit manipulations of motor effort. Third, force levels were not individually calibrated, and different participants may therefore have experienced the required force differently, potentially contributing to variability in the magnitude of the observed effects. Fourth, motor effort might have been influenced by the visual presentation used in the study, it may therefore be worthwhile for future research to dissociate the visual presentation of effort from the actual motor costs associated with executing the response, in order to determine the representational format through which motor effort is integrated into the decision process. Finally, the current design still relies on a sequential decision–action structure, rather than decisions occurring in a more naturalistic environment in which movements unfold on the fly. Future research is therefore needed to investigate whether similar effects emerge while engaging in ongoing actions.

To conclude, the present study investigated the temporal evolution of perceptual decision formation in an orientation discrimination task in which two alternative choices were reported with different levels of motor effort. In line with earlier studies, our results demonstrated that motor effort biased perceptual decisions toward the low-effort option. Motor effort influenced decision formation at multiple stages: it biased preparatory attentional allocation before stimulus presentation and later-stage evidence accumulation preceding response execution toward low-effort options, while leaving early sensory encoding of the two options unaffected. Importantly, only the modulation of later-stage evidence accumulation was associated with trial-level choice behavior, leading to an increased likelihood of selecting the low-effort option. Together, these findings suggest that motor effort primarily biases perceptual decisions during later-stage evidence accumulation preceding response execution rather than during early sensory-attentional processing. These findings contribute towards the neural basis of the embodied perceptual decision-making framework by demonstrating that motor effort can be incorporated into the decision formation process rather than merely shaping action tendencies, while providing additional temporal insights into when this effect occurs. Understanding how motor effort shapes decision formation and identifying associated neural markers can help us better understand the neural mechanisms underlying decision-making abnormalities in movement disorders, and eventually inform clinical practice and interventions.

## CRediT authorship contribution statement

**Ningjing Cui**: conceptualization, data curation, formal analysis, investigation, methodology, software, visualization, writing – original draft. **Jeroen B. J. Smeets**: conceptualization, methodology, supervision, writing – review and editing. **Robert J. van Beers**: conceptualization, methodology, supervision, writing – review and editing. **Dock H. Duncan**: conceptualization, methodology, writing – review and editing. **Bernadette C. M. van Wijk**: conceptualization, funding acquisition, methodology, supervision, writing – review and editing.

## Data availability

Behavioral data, processed EEG data, and code are available on Zenodo (link:10.5281/zenodo.23167590).

## Supplementary Materials

**Table S1.** Group-averaged psychometric parameters (threshold, width, slope, and lapse rate) for each effort-mapping condition. Values are expressed as mean ± SD across participants.

| Condition | Threshold ($^{\circ}$) | Width ($^{\circ}$) | Slope (prob/$^{\circ}$) | Lapse Rate (%) |
| --- | --- | --- | --- | --- |
| Low effort: left | 6.42 ± 5.16 | 11.48 ± 7.31 | 0.05 ± 0.04 | 0.66 ± 2.18 |
| Low effort: right | 2.83 ± 3.91 | 10.83 ± 5.78 | 0.05 ± 0.02 | 1.06 ± 1.67 |

### Supplementary Methods

To examine the potential computational mechanisms underlying the motor effort– related decision bias observed in the behavioral data, we fitted a drift-diffusion model (DDM) to each participant’s choice and RT data. The DDM is a sequential sampling model widely used to characterize binary decision-making, in which decisions are described as the gradual accumulation of noisy evidence toward one of two decision boundaries representing the available choices (Ratcliff et al., 2016; Ratcliff & McKoon, 2008). Key parameters include drift rate (*v*) representing the rate of evidence accumulation; decision boundary (*B*), representing the evidence threshold required to commit to one of the two choices; starting point (*z*) representing an initial bias toward one decision boundary prior to evidence accumulation onset; and non-decision time representing the time needed for sensory and motor processes unrelated to evidence accumulation. Models were implemented in Python 3.12.4 using the PyDDM package (Shinn et al., 2020), which employs a numerical solution to the Fokker– Planck equation, providing efficient and reliable parameter estimation at the individual level (De Gee et al., 2025; Shinn et al., 2020; White et al., 2024).

Here, we considered that motor effort-related decision bias may be reflected in changes in evidence accumulation, indexed by drift rate, and/or in shifts in the initial bias preceding evidence accumulation, indexed by starting point, in line with previous work (Ding & Gold, 2012; Hagura et al., 2017; Hanks et al., 2006, 2011; Mulder et al., 2012). For each participant, we fitted an identical model structure, selected through model selection (**Table S2** and **S3**), in which both drift rate and starting point varied as a function of motor effort. Drift rate was specified as a linear function of orientation difference, effort side, and a constant leftward choice bias:

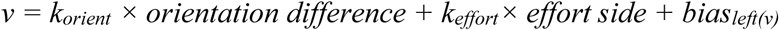

Here, *orientation difference* reflects the orientation difference of the two stimuli, and *effort side* indicates the location of the low-effort option (−1 = left; +1 = right). The *bias_left(v)_* represents a constant leftward choice bias, as indicated by the positive psychometric threshold shift observed for both effort-mapping conditions (see Section 3.1, **Figure 3B**). This parameter was incorporated into both the drift rate and starting point because its inclusion improved model fit during model selection (**Table S2** and **S3**). An example fit of the selected model for an individual participant’s RT distributions is illustrated in **Figure S1**. Positive *k_effort_* values indicated that motor effort shifted drift rate toward the low-effort option, whereas negative values indicated shifts toward the high-effort option.

The starting point (*z*) was set to vary as a function of effort side and incorporated the leftward choice bias:

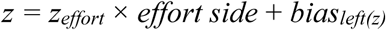

The decision boundary (B) was included as a free parameter with the upper and lower boundaries (B and −B) corresponding to right and left choices, respectively. Diffusion noise was fixed at 1 for parameter scaling, and non-decision time was modelled using a normal distribution (mu, sigma). A uniform mixture parameter was included to account for contaminated responses (e.g., random guesses) not explained by the evidence accumulation process. Positive *k_effort_* values indicated that motor effort shifted drift rate toward the low-effort option, whereas negative values indicated shifts toward the high-effort option. Positive *z_effort_* values indicated that motor effort shifted the starting point toward the low-effort option, whereas negative values indicated shifts toward the high-effort option. Plausible parameter ranges were determined based on prior DDM research and visual inspection of model fits (see **Table S4** for parameter ranges).

To validate the fitting procedure, a parameter recovery analysis (Shinn et al., 2020; Wilson & Collins, 2019) was conducted in which simulated data were generated for each participant using the obtained parameter estimates as fixed parameters while keeping the original trial-wise condition structure (i.e., the same number of trials, orientation differences and effort-mapping conditions). The model was then refitted to the simulated data, and recovered parameter estimates were compared with the true parameter values using Pearson correlations and root mean squared error (RMSE).

To examine whether effort-related model parameters were associated with the observed decision bias, we conducted separate one-sample Wilcoxon signed-rank tests for the effort effects on drift rate (*k_effort_*) and starting point (*z_effort_*) against zero. To test whether variations in these parameters explain individual choice bias, we computed between-subject Spearman correlations between participants’ decision bias and their effort-related effects on drift rate (*k_effort_*) and starting point (*z_effort_*). Non-parametric partial correlations were additionally conducted for both parameters to test whether these relationships remained significant after controlling for the other parameter.

**Table S2.** Model structures for four drift-diffusion models including a constant leftward choice bias parameter added to drift rate (*v*) and/or starting point (*z*). All models were fit to all individual participant data.

| Parameter | Model 1 | Model 2 | Model 3 | Model 4 |
| --- | --- | --- | --- | --- |
| $k_{orient}$ | ✓ | ✓ | ✓ | ✓ |
| $k_{effort}$ | ✓ | ✓ | ✓ | ✓ |
| $bias_{left(v)}$ | — | ✓ | — | ✓ |
| $bias_{left(z)}$ | — | — | ✓ | ✓ |
| B | ✓ | ✓ | ✓ | ✓ |
| $z_{effort}$ | ✓ | ✓ | ✓ | ✓ |
| mu | ✓ | ✓ | ✓ | ✓ |
| sigma | ✓ | ✓ | ✓ | ✓ |
| umixturecoef | ✓ | ✓ | ✓ | ✓ |
| Number of parameters | 7 | 8 | 8 | 9 |

**Table S3.**
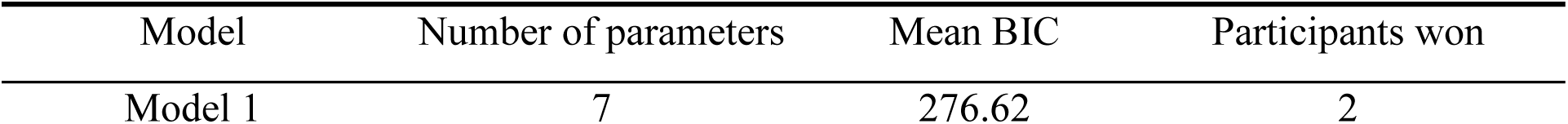
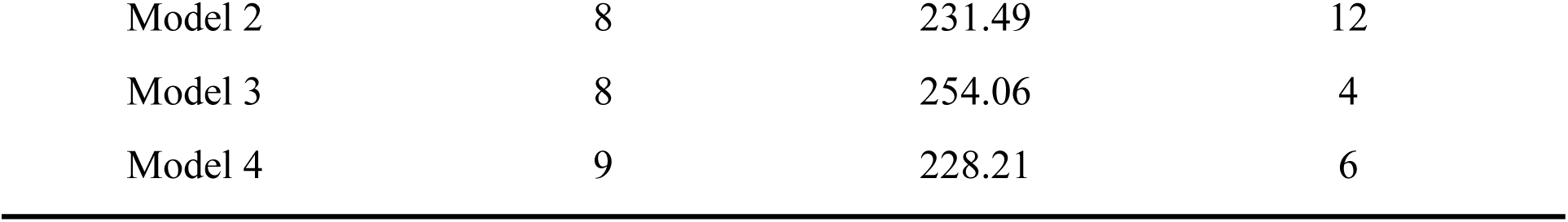
Model comparisons were performed using individual participant BIC values for each model. Model 4, which included a constant leftward choice bias parameter to both the drift rate (*v*) and starting point (*z*), yielded the lowest mean BIC across participants. Model 4 was therefore selected as the winning model and subsequently fitted to each participant’s data to obtain parameter estimates for direct comparisons with behavioral measures.

| Model | Number of parameters | Mean BIC | Participants won |
| --- | --- | --- | --- |
| Model 1 | 7 | 276.62 | 2 |
| Model 2 | 8 | 231.49 | 12 |
| Model 3 | 8 | 254.06 | 4 |
| Model 4 | 9 | 228.21 | 6 |

**Table S4.** Plausible parameter ranges for DDM fitting.

| Parameter | Range |
| --- | --- |
| $k_{orient}$ | $(-2, 2)$ |
| $k_{effort}$ | $(-3, 3)$ |
| $bias_{left(v)}$ | $(-3, 3)$ |
| $bias_{left(z)}$ | $(-0.8, 0.8)$ |
| B | $(0.3, 3)$ |
| $z_{effort}$ | (−0.8, 0.8) |
| $\mu$ | (0.1, 1) |
| $\sigma$ | (0.01, 0.15) |
| $umixturecoef$ | (0, 0.03) |

**Figure S1.**
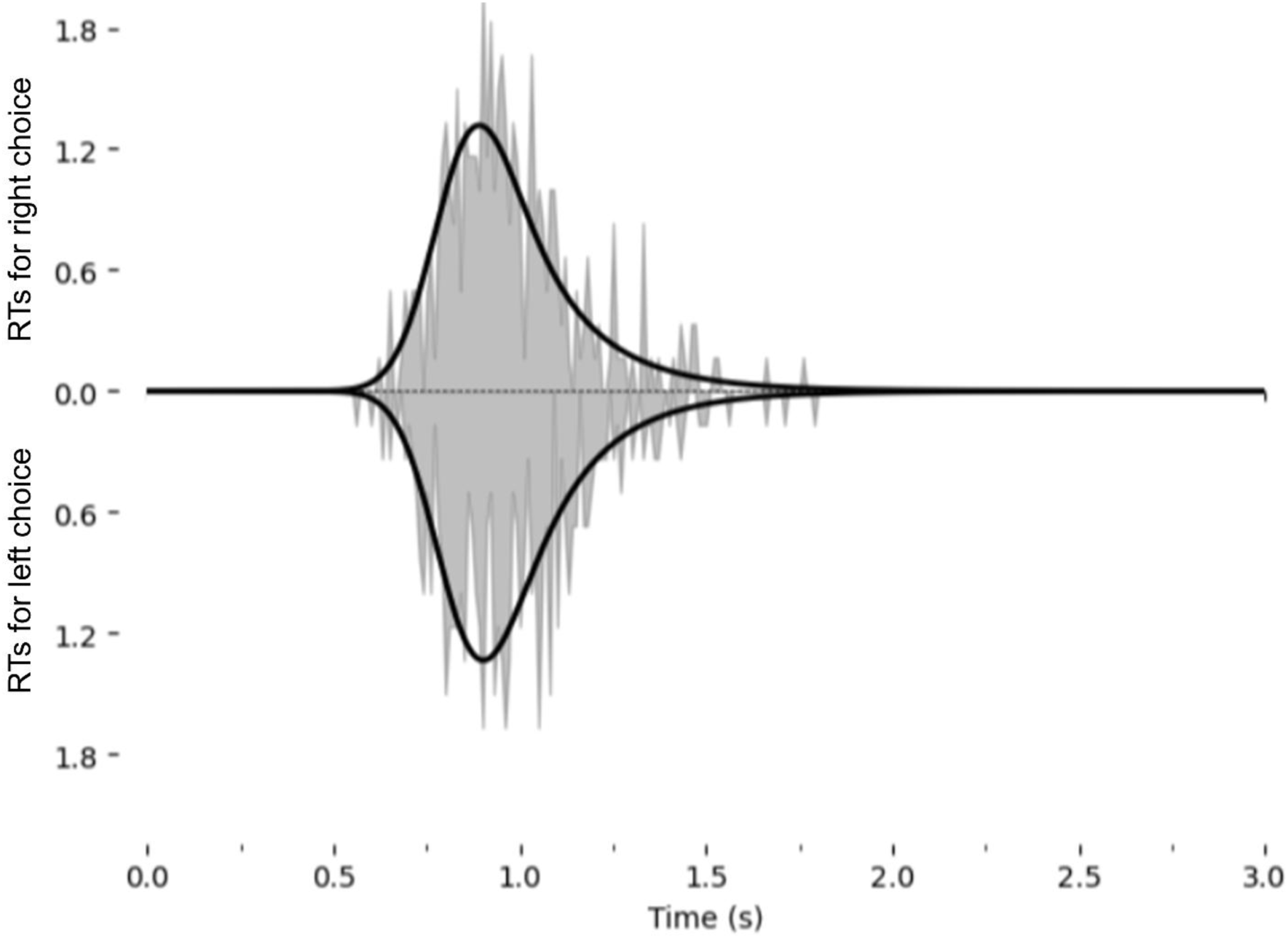
Example fit using the selected model (Model 4) to an individual participant’s RT distributions for left and right responses. The grey areas show the observed distributions, and the black curve shows the model predicted RT probabilities.

### Supplementary Results

To characterize the computational mechanisms underlying the observed motor effort-related decision bias, we fitted a DDM to each participants’ RT and choice data in which both drift rate (*v*) and starting point (*z*) varied as a function of effort mapping. As shown in **Figure S2A** for individual fitted parameter estimates, most participants exhibited positive *k_effort_* values, indicating that motor effort biased drift rate toward the low-effort option, as confirmed by the significant positive effect (median = 0.34, *z* = 3.8, *p* < .001). By contrast, most participants exhibited negative *z_effort_* values, suggesting a weak starting point bias toward the high-effort option (median = –0.1, *z* = -2.77, *p* = .006), suggesting a potential trade-off between the two parameters. A parameter recovery analysis was conducted by simulating data using the same number of trials and trial-wise condition structure as the original dataset while fixing parameter values to each participant’s fitted parameter estimates. The model was then refitted to the simulated data, and recovered parameter estimates were compared with the true estimates. As shown in **Figure S2B**, all parameters showed strong recovery (all r ≥ 0.8, RMSE < 0.25), suggesting reliable parameter estimation. To further examine whether variations in the drift rate or starting-point bias explained the individual behavioral choice bias, Spearman correlations were computed between behavioral decision bias, quantified as the psychometric threshold shift between effort-mapping conditions, and both *k_effort_* and *z_effort_* values. As shown in **Figure S2C**, decision bias was significantly correlated with *k_effort_* (rho = 0.56, *p* = .005) but not with *z_effort_* (rho = 0.31, *p* = .141). Considering the potential shared variance between the two parameters, as indicated by their significant correlation (rho = –0.50, *p* = .015), we additionally conducted partial correlations to test whether each parameter independently correlated with decision bias while controlling for the other parameter. Both *k_effort_* and *z_effort_* were significantly associated with decision bias after controlling for the other parameter (*k_effort_* controlling for *z_effort_*: rho = 0.87, *p* < .001; *z_effort_* controlling for *k_effort_*: rho = 0.82, *p* <.001). These findings indicate that motor effort-related decision bias may be reflected in both drift-rate and starting-point parameters.

**Figure S2.**
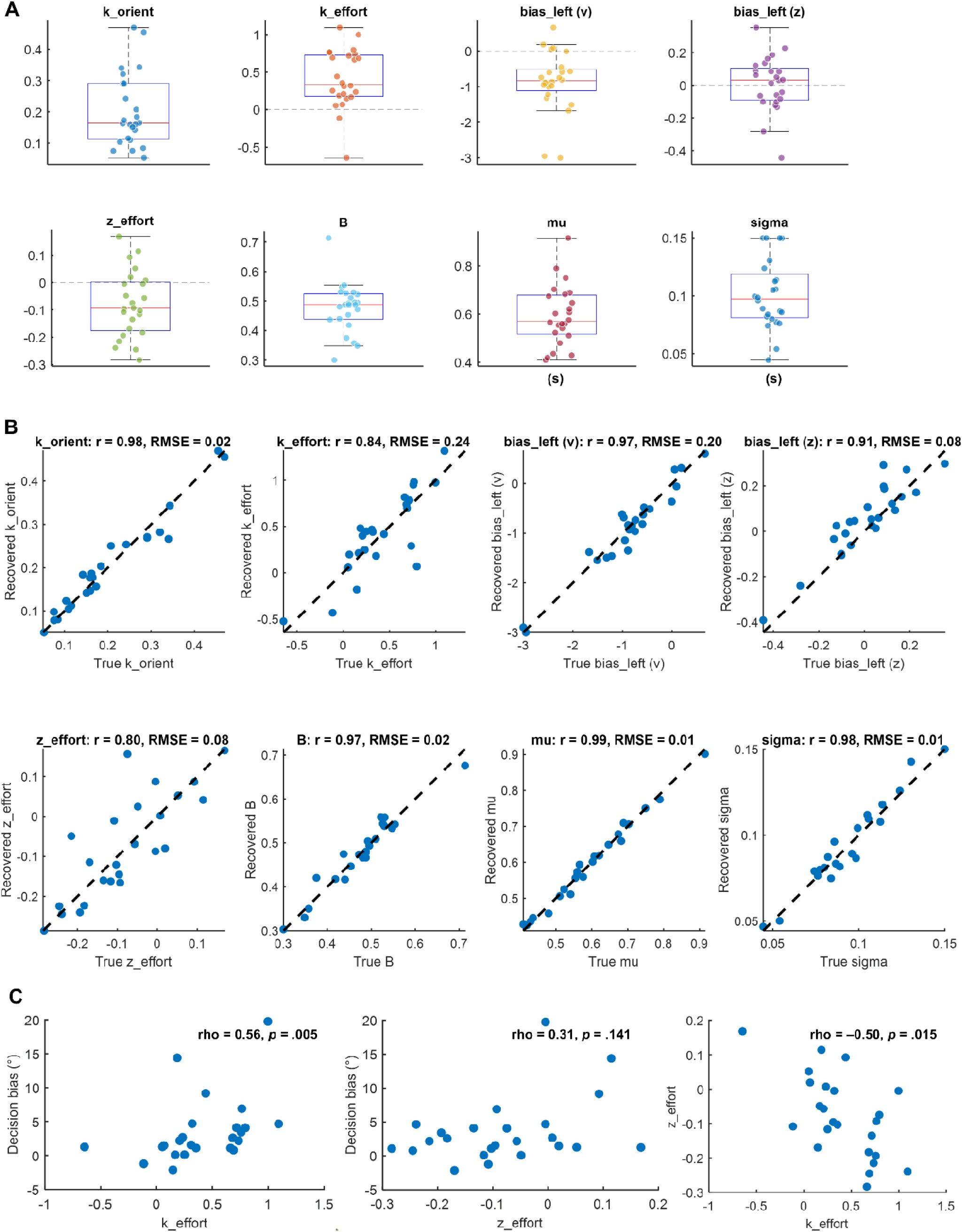
**(A)** DDM parameter estimates for individual participants after fitting a model in which drift rate (v) was specified as a linear function of orientation difference, effort side, and a constant leftward choice bias: *v = k_orient_ × orientation difference + k_effort_× effort side + bias_left(v)_.* The starting point (*z*) was set to vary as a function of the effort side with a constant leftward choice bias: *z = z_effort_ × effort side + bias_left(z)_.* The decision boundary (B) was included as a free parameter with the upper and lower boundaries corresponding to right and left choices, respectively. Diffusion noise was fixed at 1 for parameter scaling, and non-decision time was modelled using a normal distribution (mu, sigma). A uniform mixture parameter was included to account for contaminated responses (e.g., random guesses) not explained by the evidence accumulation process. **(B)** Parameter recovery analysis showing strong correlations between recovered and true parameter estimates (all r > 0.8, RMSE < 0.2). **(C)** A significant correlation was observed between decision bias and *k_effort_* (*p* = .005) but not *z_effort_* (*p* = .141), though a negative correlation was found between *k_effort_* and *z_effort_* (*p* = .015). Each blue dot represents an individual participant.

